# Dual roles of IQCG, a novel microtubule nucleation factor rapidly evolving in humans

**DOI:** 10.1101/2024.12.16.628806

**Authors:** Xingtang Liu, Yongzhen Wu, Xia He, Yuan Shen, Wenxiang Fu, Haiyang Hu, Haijing Yu

**Affiliations:** School of life sciences, Yunnan University, Kunming, China; Center for life sciences, Yunnan University, Kunming, China; Central Laboratory, Shanghai Pulmonary Hospital, Tongji University School of Medicine, Shanghai, China; Yunnan International Joint Laboratory of Virology & Immunology, Yunnan University, Kunming, China

**Keywords:** IQCG, centrosome, microtubule nucleation factor, cell division, rapid evolution

## Abstract

The centrosome serves as the main microtubule-organizing structure, playing crucial roles in animal development. Uncovering new components and elucidating their functions helps to understand the regulation of cell maintenance and division by the centrosome. In this study, through a combination of genetic analyses and cell biological characterizations, we identify IQ Motif Containing G (IQCG) as a novel microtubule nucleation factor with a dual role involved in both centrosomal microtubule organization and mitotic spindle formation. IQCG interacts with GSK3β via its N2 region, enhancing its centrosomal accumulation in interphase; subsequently, IQCG disperses into the spindle to promote microtubule generation during mitosis and ensure robust cell division. Furthermore, our analyses indicate that the *IQCG* gene has undergone accelerated evolution within the human lineage, with positive selection acting on the variable N2 region. This adaptation may confer advantages to human cells by augmenting central regulatory control over cellular activities through the centrosome. Our findings offer novel insights into the functional mechanisms and evolutionary significance of the human centrosome associated with human evolution.

**Summary blurb:** The cellular functions of the novel microtubule nucleation factor IQCG, which enhances centrosomal microtubule organization and cell division robustness, are implicated in human evolution.

## Introduction

Centrosome, the major microtubule organizing center in animal cells, is involved in regulations on cell division, motility and signaling (Wu and Akhmanova 2017; Vasquez-Limeta and Loncarek 2021). The canonical centrosome comprises two centrioles embedded within pericentriolar matrix (PCM). This organelle is an ancient structure with a complex evolutionary history, displaying both conserved and divergent features across species (Bornens and Azimzadeh 2007; Ito and Bettencourt-Dias 2018). Novel modifications in centrosomal components and their functions have evolved alongside the diversification of organisms (Ito and Bettencourt-Dias 2018; Hodges et al. 2010). A significant increase in the number of centrosomal genes from *Drosophila* to humans has been documented (Alves-Cruzeiro et al. 2014), and proteins appear to have been incrementally added to the centriole structure in a stepwise manner throughout evolution (Carvalho-Santos et al. 2010; Zheng et al. 2016). The complexity of disordered and coiled-coil regions in centrosome proteins was indicated to correlate with an increase in cell type diversity, contributing to greater animal complexity (Nido et al. 2012). In contrast to centrioles, proteins found in the PCM are responsible for microtubule nucleation, modification, and stability; this empowers the centrosome to function as an microtubule organization center (MTOC), thereby regulating both the quantity and composition of cellular microtubule cytoskeletons (Woodruff et al. 2014). Variations in size and organizational capacity of the PCM across species reflect pronounced evolutionary adaptations tailored to functional requirements during development (Magescas et al. 2021; Lee et al. 2021).

In humans, mutations underline the necessity of centrosomal functions for proper neocortex formation (Wilsch-Bräuninger and Huttner 2021). Distinct selective signatures are evident among genes associated with human primary microcephaly—characterized by a reduced cerebral cortex—indicating that evolutionarily advantageous modifications in centrosomal functions are closely linked to the expansion of the human neocortex (Xu et al. 2017; Montgomery et al. 2011; Ponting and Jackson 2005; Gilmore and Walsh 2013). A deeper understanding of innovations related to the centrosome could illuminate biological mechanisms underlying human evolution.

The *IQCG* gene is one of the target genes of the human-specific microRNA miR-941 that we previously reported (Hu et al. 2012). *IQCG* encodes an IQ motif-containing protein implicated in male infertility and several types of cancer (Zhu et al. 2018; Pan et al. 2008). Studies have suggested that *IQCG* and its orthologues are involved in flagellum and microtubule functions (Bower et al. 2013; Harris et al. 2014; Li et al. 2014). In mice, *IQCG* is essential for spermatogenesis, where it is highly expressed and co-localizes with a transient skirt-like cytoskeletal structure manchette, and depletion of *IQCG* results in male infertility (Harris et al. 2014; Li et al. 2014). In zebrafish development, *IQCG*-deficient embryos are lack of mitotic haematopoietic stem cells (Chen et al. 2014). However, the cellular roles of *IQCG* remain to be fully clarified.

In this study, we demonstrate that the *IQCG* gene encodes a novel microtubule nucleation factor and has undergone rapid evolution in the human lineage. Characterized by strong positive selection on the N2 region of the human IQCG protein, it may have acquired advanced functionalities to enhance robustness in human cell division. This study offers novel insights into human centrosome organization and potential adaptive modifications pertaining to its functions.

## Results

### The human *IQCG* gene encodes a centrosome protein

We first carried co-expression analyses on gene expression data from 53 human tissues in the GTEx database (GTEx Consortium 2013) to predict the involvement of *IQCG* in biological processes. The results indicated that it primarily participates in cilium/flagellum organization and microtubule functions (Table S6), which aligned with its previously reported roles (Harris et al. 2014; Li et al. 2014; Chen et al. 2014; Bower et al. 2013). To investigate its cellular function, we examined the subcellular localization of endogenous IQCG protein through immunostaining using a IQCG-specific antibody, which revealed a cytoplasmic distribution with a significant portion localized at the centrosome (Fig 1A, left). This centrosome localization was further confirmed by observing ectopically expressed eGFP-tagged IQCG (Fig 1A, middle and right). Importantly, this distribution pattern remained unchanged even after disrupting microtubules with Nocodazole (Fig 1B), indicating that IQCG localizes to the centrosome independently of the microtubule network.

**Figure 1.**
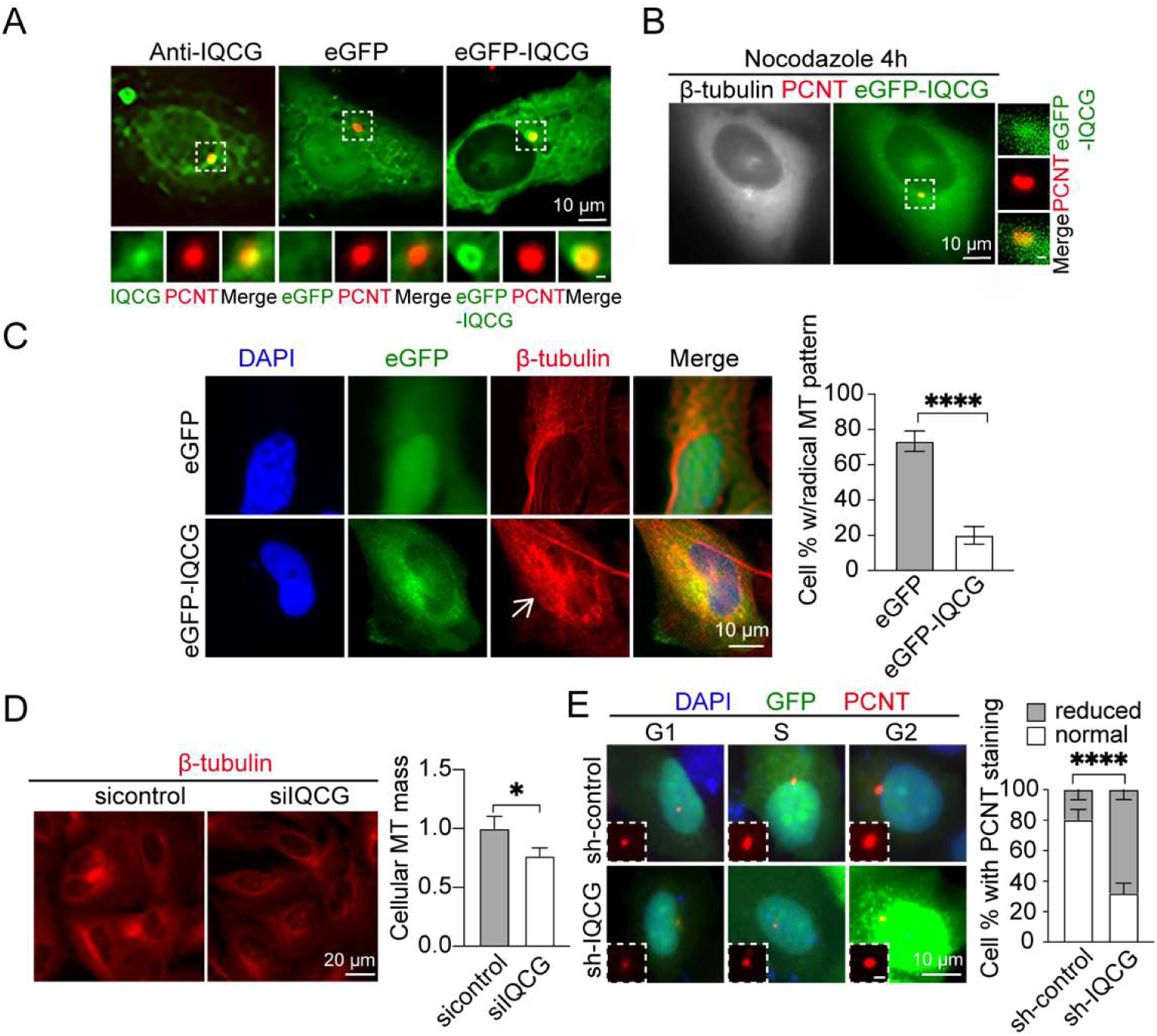
The human IQCG protein localizes to the centrosome and regulates microtubule organization. **(A)** Immunofluorescence staining reveals endogenous IQCG localization at the centrosome (left), while transiently expressed eGFP-IQCG confirms this localization (right) in HeLa cells. EGFP alone serves as a control (middle). **(B)** Following Nocodazole treatment to disassemble microtubules, IQCG remains localized at the centrosome. Boxed areas are enlarged for detail. **(C)** Expression of IQCG alters the distribution of microtubules, and analysis of cell proportion with typical radial MT distribution pattern from the centrosome is performed in control and IQCG-expressing HeLa cells. An arrow indicates altered MT distribution. n = 200. **(D)** Knockdown of IQCG by siRNA targeting IQCG decreases total cellular microtubule mass revealed by immunostaining in HeLa cells, and reduced cellular MT intensity is analyzed in control and IQCG-knockdown cells. n = 1000. **(E)** Knockdown of IQCG results in reduced recruitment of PCNT at the centrosome during G1, S, and G2 phases in HeLa cells transfected with shRNA targeting IQCG. Insets provide magnified details. Green fluorescence marks shRNA expression. Analysis is conducted on cell proportions exhibiting normal or reduced centrosomal PCNT staining in control or IQCG-shRNA expressing cells. n = 200. Scale bars: 10 or 20 μM as indicated within images, 1 μM for insets. Font color corresponds to respective fluorescence stainings observed within images. Blue nucleus is stained with DAPI. \**p* < 0.05, \*\*\*\**p* < 0 .0001 At least three independent experimental replications were performed.

Overexpression of IQCG led to altered MT patterns showing enhancement of tubulin signals where IQCG was presented, in contrast to the typical MT pattern with radial arrays emanating from the centrosome in control cells (Fig 1C). Conversely, IQCG depletion using siRNA led to a noticeable reduction in cellular MT mass with a disrupted radical pattern (Fig 1D and Fig S1A), which was further supported by knockdown experiments with shRNA (Fig S1B). In *IQCG* knockdown cells, we also observed decreased PCNT staining at centrosomes throughout interphase (Fig 1E). This reduced staining was accompanied with diminished recruitment or vague staining of other critical centrosome proteins such as Ninein, Rootletin and PCM1 (Pericentriolar Matrix 1), as well as the MT minus-end protein CAMSAP2 (Calmodulin Regulated Spectrin Associated Protein Family Member 2) (Fig S1C), suggesting the integrity of centrosomal organization was affected by IQCG. These findings underscored the important roles played by IQCG in microtubule and centrosome organization.

### The human IQCG serves as an MT nucleation factor in PCM

To determine the exact function of IQCG in the centrosome, we first used a series of well-established centrosome proteins as landmarks to ascertain the precise position of the IQCG protein within the architecture of the centrosome. The results from immunostaining confocal microscopy revealed that the spatial distribution of IQCG overlapped with that of PCM scaffold protein PCNT, albeit being more diffuse. In contrast, no co-localization was observed with centrioles or centriolar satellites represented by dotted Centrin and punctate PCM1 staining respectively, indicating its specific positioning in the outer layer of PCM (Fig 2A).

**Figure 2.**
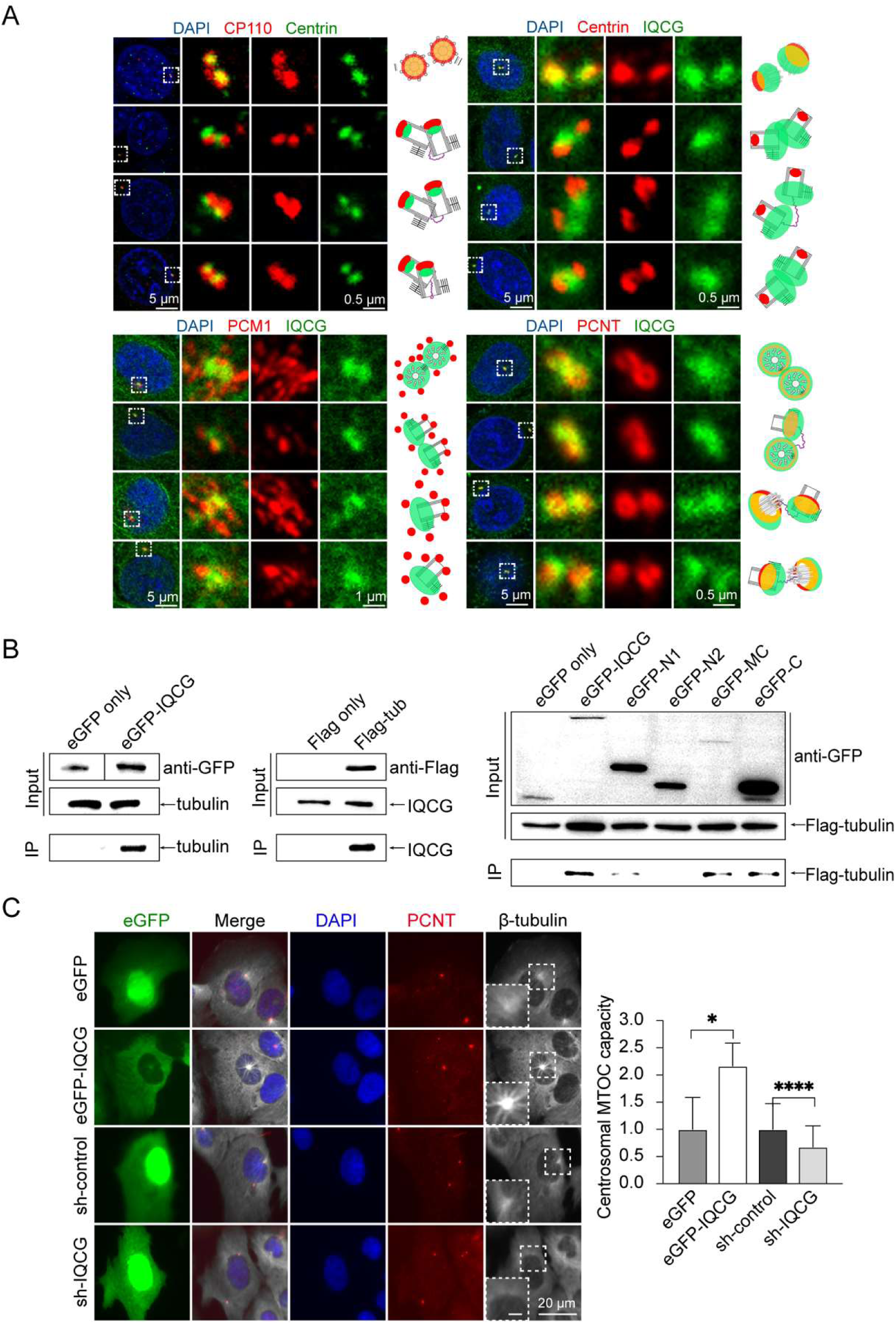
The IQCG protein localizes specifically in the outer layer of the PCM within the centrosome and interacts with tubulin via its C region. **(A)** Immunofluorescence staining using antibodies reveals that IQCG is present in the PCM outer layer of the centrosome, overlapping with PCNT. Distal regions of the centrioles are stained with Centrin (green) and CP110 (red), and accurate localization of IQCG (green) in relation to Centrin, PCNT, and PCM1 (red) is observed in HeLa cells. Schematic representations illustrate their relative positions within the centrosome. Scale bars: 5 µm for whole cell images; 0.5 µm or 1 µm for enlarged images. **(B)** Co-immunoprecipitation assays demonstrate that eGFP-IQCG precipitates tubulin but not eGFP alone; reciprocally, Flag-tubulin precipitates IQCG but not Flag alone; and the interaction between IQCG and tubulin is mediated via its C region. Lysates from HEK293T cells expressing bait proteins (eGFP only, eGFP-IQCG and its fragments, Flag only or Flag-tubulin) are used as inputs, and immunoprecipitates (IPs) are obtained using anti-GFP or anti-Flag beads. Inputs and IPs are probed using anti-GFP, anti-Flag, anti-tubulin or anti-IQCG antibodies. **(C)** Centrosomal MTOC capacity is enhanced by IQCG expression and reduced by IQCG knockdown in U2OS cells. Insets show magnification of boxed areas. Scale bars: 10 µm for whole cell images; 10 µm for insets. Green fluorescence: eGFP in eGFP and eGFP-IQCG expressing cells, or GFP marking shRNA expression in control and *IQCG*-knockdown cells. Intensity analysis is performed on regrown microtubules at the centrosome after a 5-minute recovery following Nocodazole treatment. n = 50. At least three independent experimental replications were performed.

Next, through a co-immunoprecipitation experiment employing eGFP affinity beads, we successfully identified tubulin protein as a binding partner for eGFP-tagged IQCG; this interaction was further validated using expressed Flag-tubulin as bait in a reciprocal experiment (Fig 2B). To characterize the nature of interaction between IQCG and tubulin, MT co-sedimentation assays were conducted under conditions where MT stability was altered. By treating cells with Nocodazole or Docetaxol to depolymerize or stabilize MTs respectively, we investigated whether IQCG sedimented along with polymerized MTs. SDS-PAGE analysis showed no change in IQCG levels in soluble (S) fraction containing proteins not associated with polymerized MTs versus precipitated fraction as pellet (P) containing proteins associated with polymerized MTs, regardless of MT polymerization status (Fig S2). This result effectively established that IQCG did not associate with polymerized tubulins but rather interacted with tubulin specifically at minus ends of MTs.

Through a microtubule regrowth assay, we observed enhanced centrosomal MTOC capacity in cells expressing IQCG, as manifested by significantly larger MT asters at the centrosome following Nocodazole treatment and release; conversely, depletion of IQCG resulted in impaired MT regrowth (Fig 2C). Based on these findings, we concluded that the IQCG gene encoded a PCM component serving as an essential factor for MT nucleation at the centrosome.

### The human IQCG regulates mitotic spindle assembly

Interestingly, the cellular distribution of IQCG became dynamic upon the onset of mitosis. Specifically, IQCG no longer concentrated at the centrosome during prometaphase but dispersed to the spindle (Fig 3A). As mitosis progressed into metaphase, IQCG clearly co-localized with the spindle around the spindle pole (Fig 3B). This observed dynamic shift in IQCG localization coincided with rapid assembly of the spindle during prophase and metaphase. In anaphase, this co-localization pattern began to blur and ultimately disappeared (Fig 3B). The transfer of IQCG to the spindle is analogous to that observed for mouse IQCG transferring to the manchette at step 9 of sperm development, which also coincides with rapid manchette assembly (Li et al. 2014).

**Figure 3.**
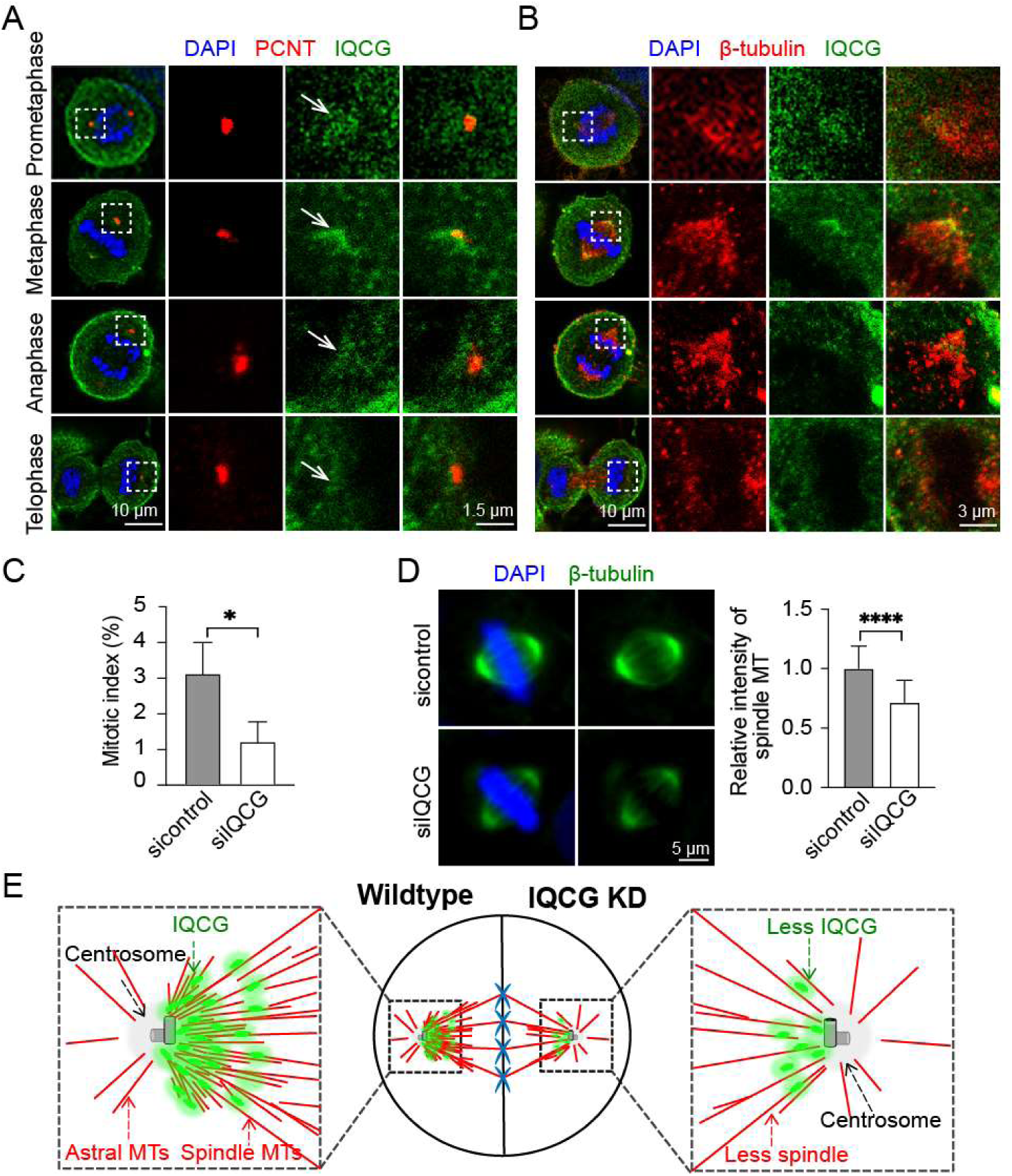
IQCG participates in mitosis by promoting spindle formation. **(A)** The concentration of IQCG at the centrosome is no longer observed in mitotic HeLa cells. Boxed areas are enlarged for detail. Arrows indicate the position of the centrosome. Blue: chromosome stained by DAPI; green: endogenous IQCG stained by anti-IQCG; red: centrosome marked by anti-PCNT. Scale bars: 10 μM for whole cells, 1.5 μM for enlarged images. **(B)** IQCG colocalizes with the spindle at the spindle pole periphery. Boxed areas are enlarged for detail. Blue: chromosome stained by DAPI; green: endogenous IQCG stained by anti-IQCG; red: MT stained by anti- β-tubulin. Scale bars: 10 μM for whole cells, 3 μM for enlarged images. **(C)** Knockdown of IQCG leads to a decrease in the mitotic index of HeLa cells. n ≥ 6000. **(D)** Depletion of IQCG results in reduced intensity of microtubules within the metaphase spindle in HeLa cells. Scale bar: 5 μM. MT intensity within the spindle is analyzed. n ≥ 30. In **(A-D)**, more than three independent experimental replications were performed. \**p* < 0.05, \*\*\*\**p* < 0.0001. **(E)** Illustrative depiction shows IQCG’s role in promoting spindle microtubule formation during mitosis.

To elucidate IQCG’s role in mitosis, we assessed mitotic phenotypes in HeLa cells following knockdown of IQCG. The mitotic index in these cells significantly decreased (Fig 3C), indicating a substantial impairment in cell division due to lack of IQCG expression. Consistent with this finding, zebrafish embryos deficient in IQCG were also found to lack mitotic stem cells marked by phosphorylated Histone 3 during hematopoiesis (Chen et al. 2014). In limited instances where mitosis took place in the IQCG-knockdown cells, their spindles appeared morphologically normal— maintaining bipolarity as well as appropriate positions and shapes—when compared with control cells. However, there was a notable reduction in microtubule density within the metaphase spindles; they presented a relatively "hollow" appearance (Fig 3D), suggesting that insufficient levels of IQCG led to inadequate microtubule production within spindles despite minimal initiation of mitosis occurring overall.

These observations implied that IQCG played a critical role in regulating both mitotic initiation and microtubule generation within spindles. Given that both spindles and manchettes are transient and dynamic microtubule structures requiring rapid nucleation processes, we concluded that a dual role for IQCG was implicated in facilitating swift assembly of these microtubule structures independently from centrosomal influence, as illustrated in Fig 3E.

### IQCG’s M and C regions are responsible for centrosome localization and MT nucleation, respectively

To investigate the functional mode of IQCG’s dual roles, we analyzed the functional domains within the human IQCG protein. This protein comprises 443 amino acids and contains several disordered regions, two coiled-coil domains, and a C-terminal IQ motif annotated by Uniprot (UniProt Consortium 2023) and InterPro (Paysan-Lafosse et al. 2023). Based on domain information and tertiary structure prediction, we divided the IQCG protein into four regions: N1 (1– 80 aa), N2 (81–149 aa), M (150–383 aa) and C (384–443 aa) for further analyses (Fig S3A). Interestingly, distinct acid-base properties were identified in these regions (Fig S3B). The N1 region was predominantly acidic, while the C-terminal region was notably basic. The N2 and M regions were approximately neutral with a rich yet balanced composition of acidic and basic amino acids. It is worth mentioning that bipartite residual regions rich in both acidic and basic amino acids were frequently seen in MT-associated proteins, which could facilitate their interaction with the MT surface. These unique compositional characteristics signified the distinctive biological roles played by these specific regions in the human IQCG protein.

To assess the functionalities of the four regions, we generated and examined a series of eGFP fusion proteins encompassing IQCG fragments: N1, N2, M, C, ΔN1, ΔN2, ΔM, ΔC, and N2M regions (Fig 4A and Fig S3C). Subcellular localization studies in HeLa cells revealed that the M region and the proteins containing it consistently targeted to the centrosome, whereas removal of the M region (ΔM) completely abolished centrosomal targeting (Fig 4A and B), indicating that solely the M region was sufficient for association with the centrosome. Intriguingly, addition of N2 to the M region (N2M) further strengthened M region-determined centrosome association (Fig 4A).

**Figure 4.**
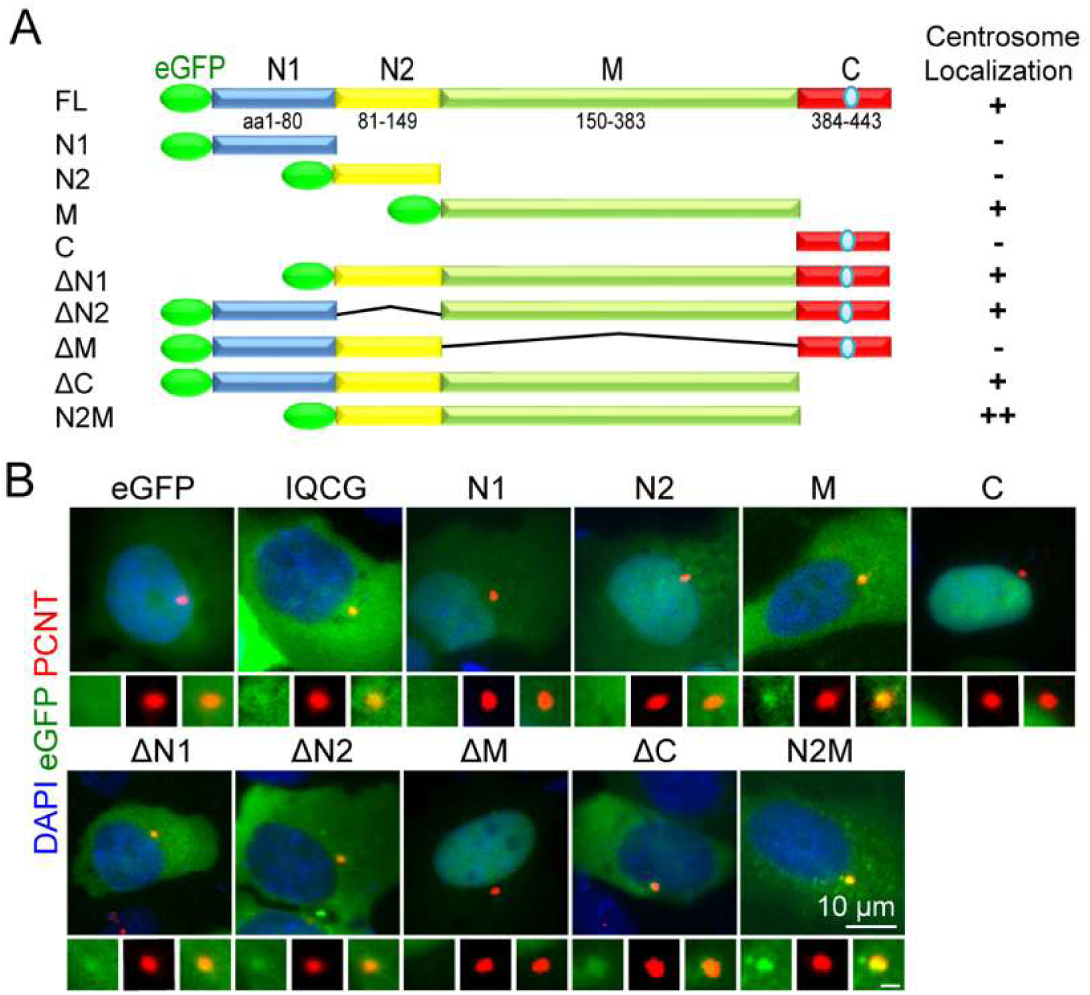
The localization of IQCG at the centrosome is determined by its M region, and this localization is further enhanced by the addition of the N2 region. **(A)** Schematic representation of chimeric eGFP-IQCG and its fragment proteins used in this study, illustrating their centrosome localization. The M-determined centrosome localization is strengthened by the presence of the N2 region. **(B)** Fluorescence microscopy images show centrosome localization of eGFP-IQCG and its fragment proteins in HeLa cells. Centrosomes are magnified for better visualization. Blue: DAPI-stained nucleus; green: fluorescence from eGFP or eGFP-fused proteins; red: centrosome marked by anti-PCNT. Scale bars: 10 µm; magnified images, 1 µm.

With co-immunoprecipitation analyses using eGFP-tagged IQCG and fragmented proteins of IQCG as bait, we discovered that primarily it was the C region, which contained an IQ motif and had a high content of basic amino acids, responsible for the interaction of IQCG with tubulin (Fig 2B). This observation suggested a crucial role for the C region in mediating IQCG’s function as an MT nucleation factor. Due to its low expression level (Fig S3C), we did not directly employ only the M region itself in the co-immunoprecipitation experiments. In summary, the M and C regions played pivotal roles in mediating IQCG’s function as an MT nucleation factor in the centrosome.

### IQCG has evolved rapidly in humans, and its positively selected N2 region mediates GSK3β interaction

As one of the target genes of the human-unique microRNA miR-941, which originated *de novo* in humans (Hu et al. 2012), we analyzed the evolutionary characteristics of the *IQCG* gene based on eleven representative primate species, including five apes (human, chimpanzee, gorilla, orangutan, gibbon), four Old World monkeys (golden snub-nosed monkey, Vervet monkey, macaca, olive baboon), and two New World monkeys (night monkey and marmoset). The rate of evolution of *IQCG*, as measured by the ratio of non-synonymous to synonymous substitutions (Ka/Ks), was significantly accelerated in the human lineage following divergence from chimpanzees (Fig 5A), indicating rapid evolutionary changes specific to humans. A sliding window-based analysis revealed that this accelerated evolution was uneven across the coding region of human *IQCG*, with a particularly prominent Ka/Ks ratio exceeding 2 within the N2 region (Fig 5B), suggesting strong positive selection acting upon this region in response to adaptive pressures specific to humans.

**Figure 5.**
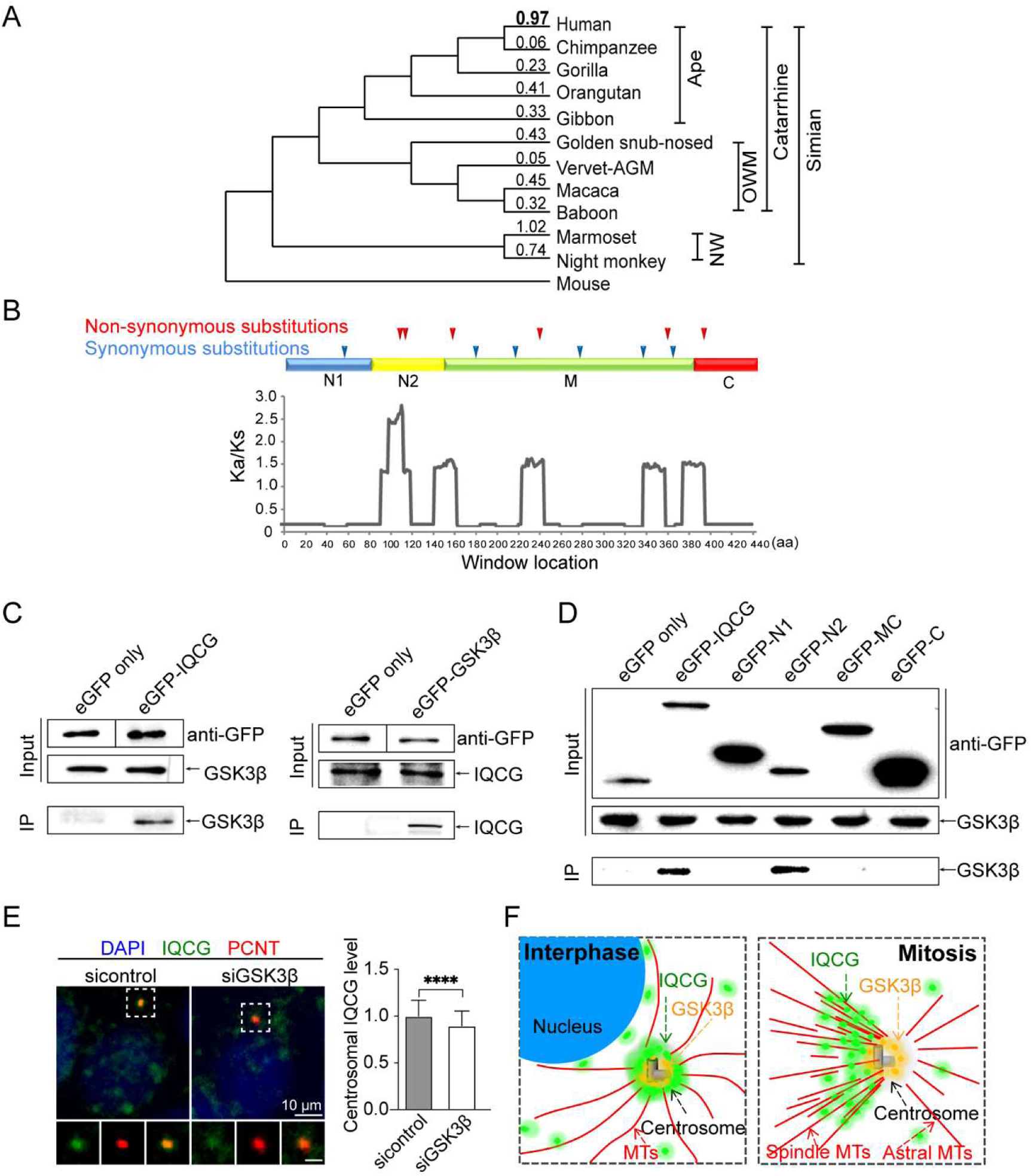
The human *IQCG* gene has undergone rapid evolution in the human lineage, with its N2 region exhibiting significant positive selection signals. This N2 region mediates the interaction of IQCG with GSK3β, enhancing its centrosome localization. **(A)** Evolutionary analysis of primates reveals evident rapid evolution of the *IQCG* gene in the human lineage, as indicated by Ka/Ks ratios above each branch of the phylogenetic tree. Branch lengths are arbitrary and do not reflect evolutionary time. OWM: Old World Monkey; NWM: New World monkey. **(B)** Sliding-window analysis reveals uneven positive selection pressure along the coding region of human *IQCG* (annotated with amino acid position), where significent positive selection signals are identified within the N2 region. Peaks above the threshold (dotted line, Ka/Ks = 1) indicate an excess of non-synonymous substitutions over neutral selection. The positions of mutations are indicated above. **(C)** Co-immunoprecipitation assays demonstrate reciprocal interaction between IQCG and GSK3β. GSK3β is precipitated with eGFP-IQCG but not with eGFP alone, and IQCG is precipitated with eGFP-GSK3β but not with eGFP alone. **(D)** The N2 region of IQCG mediates its interaction with GSK3β. In **(C-D)**, inputs are lysates of HEK293T cells expressing eGFP only or eGFP-bait proteins (IQCG, its fragments, or GSK3β). IPs are obtained using anti-GFP beads. Inputs and IPs are probed using anti-GFP, anti-IQCG or anti-GSK3β antibodies. **(E)** Knockdown of GSK3β impairs centrosome localization of IQCG in HeLa cells. Boxed areas are magnified. Green: endogenous IQCG detected by anti-IQCG; blue: nucleus stained with DAPI; red: centrosome marked by anti-PCNT. Scale bars: 10 µm for whole cells; 1 µm for magnified images. Centrosomal levels of IQCG revealed by immunostaining were evaluated in control and GSK3β-knockdown cells. n≥300. \*\*\*\**p* < 0.0001. In **(C-E)**, at least three independent experimental replications were performed. **(F)** Illustrative depiction highlights that during interphase, the interaction between IQGC and GKS3β enhances its centrosome localization; whereas in mitosis, IQCG dissociates from GSK3β at the centrosome to fulfill its dual role in promoting spindle formation.

Through co-immunoprecipitation experiments (Fig 5C) and protein-protein interaction databases as described in Methods section, glycogen synthase kinase-3β (GSK3β) was identified as a common interacting partner for IQCG. Furthermore, it was found that the interaction between IQCG and GSK3β was mediated by the N2 region of IQCG (Fig 5D). Subcellular localization studies using immunostaining and eGFP-fused protein expression demonstrated that GSK3β also prominently localized at the centrosome (Fig S4A and B). Silencing *GSK3β* by siRNA resulted in a slight but noticeable reduction in centrosomal levels of IQCG (Fig 5E and Fig S4C), providing support for its role in promoting centrosome recruitment of IQCG. This finding also explained enhanced centrosome association observed for IQCG-N2M since N2 interacted with GSK3β and further recruited IQCG to the centrosome.

However, our investigation into the relationship between IQCG and GSK3β during mitosis revealed that despite IQCG translocating to the spindle, GSK3β remained localized in the centrosome throughout mitosis as indicated by immunostaining (Fig S4D). This observation suggested that the interaction between IQCG and GSK3β was limited to interphase, indicating that IQCG’s function during mitosis did not directly involve GSK3β. We postulated that the interaction between IQCG-N2 and GSK3β facilitated the centrosomal accumulation of IQCG during interphase, subsequently enhancing efficient spindle assembly during mitosis and contributing to robust cell division (Fig 5F).

Accordingly, based on the protein sequence alignment of IQCG orthologs from ten vertebrates, we identified high levels of amino acid substitution in the N2 region starting from zebrafish and persisting among six mammals (Fig S5). This observation demonstrated a high adaptability of the N2 region which has undergone progressive evolution, particularly evident in humans with a strong selective signal as mentioned earlier. Overall, these results supported an adaptive role of the N2 region in animal evolution which may have acted on cell function to enhance robustness of cell division.

In summary, our study revealed that the *IQCG* gene encoded a novel microtubule nucleation factor playing a dual role in centrosomal MTOC and spindle formation. The fast-evolving N2 region in the human IQCG protein enhanced robustness of cell division by facilitating adequate IQCG loading at the centrosome via its interaction with GSK3β. Our findings provided new insights into the functional mechanisms and evolutionary implications associated with the human centrosome.

## Discussion

In this study, we discovered that the *IQCG* gene encoded a novel MT nucleation factor critical for centrosomal MT organization and cell division. The IQCG-interacting protein GSK3β is a multi-tasking kinase involved in regulating various aspects of cellular physiology, particularly playing significant signaling roles in brain development (Hajka et al. 2021; Luo 2012). Although the centrosomal role of GSK3β remains unclear, its involvement in mitotic entry and spindle assembly has been observed (Dewi et al. 2018; Lee et al. 2013). Our findings on the essential role of IQCG in mitosis suggested that the interaction between GSK3β and IQCG may contribute to the mitotic function of GSK3β. Furthermore, Ninein, initially identified as a GSK3β-interacting protein (Hong et al. 2000), localizes at the mother centriolar appendages and also serves as an MT nucleating and anchoring protein (Mogensen et al. 2000). From a functional perspective, it is plausible that IQCG, GSK3β, and Ninein form a functional complex synergistically cooperating in MT organization.

The mitotic spindle plays a critical role in segregating genetic material into daughter cells during cell division (Kraus et al. 2023). Efficient spindle assembly relies on multiple nucleation pathways including centrosome-, chromosome-, kinetochore- and MT-mediated nucleation to rapidly generate thousands of microtubules (Valdez et al. 2023). In human cells, MT-mediated branching nucleation generates the majority of spindle microtubules, but the amplification of spindle MTs was impaired by the depletion of IQCG in this study. Considering mouse IQCG’s involvement in manchette assembly where other pathways of MT nucleation from centrosomes, chromosomes or kinetochores are absent, we speculated that IQCG actively participated in MT-mediated MT nucleation. Nonetheless, further investigation is required to elucidate the detailed mechanism underlying IQCG’s contribution to MT-mediated MT nucleation.

Over time, there has been an evolutionary trend in the concentration of components at the centrosome (Alves-Cruzeiro et al. 2014; Carvalho-Santos et al. 2010). For example, Ninein localizes to the appendages of the mother centriole in mammals (Mogensen et al. 2000), whereas its *Drosophila* homologous protein is found at the periphery of the centrosome (Zheng et al. 2016). Similarly, increased recruitment of IQCG to the centrosome through its variable N2 region represents a comparable scenario. This results in enhanced MTOC capability and reinforces central control over cellular activity mediated by the centrosome. Moreover, maintaining cell division potential heavily relies on the MTOC capability of the centrosome since its attenuation can be coupled to cell differentiation (Sanchez and Feldman 2017). It has also been observed that older centrosomes exhibiting higher MTOC activity are preferentially present in renewing progenitor cells, while daughter centrosomes are selectively inherited by differentiating progeny (Yang et al. 2021). Therefore, from an evolutionary perspective, progressive gain in MTOC capability within the centrosome, exemplified by the case of IQCG, may have accompanied organismal evolution and contributed to enhanced cell division potential in humans.

Lastly, the *IQCG* gene is among the target genes of miR-941, a microRNA specific to humans that we previously reported on and which may have played a significant role in shaping human evolution (Hu et al. 2012). This study establishes a crucial link between the regulatory network of miR-941 and the intricate functions of centrosomes associated with human evolution. Our investigation into the function of the *IQCG* gene highlighted its canonical roles in human cell lines, while its functions in specific cellular contexts, such as neural cells, remain to be elucidated. Although no studies have yet linked *IQCG* with human brain development thus far, its recognition in Diamond-Blackfan anemia (DBA)—a disorder characterized by bone marrow failure—is noteworthy. Notably, *IQCG* is located adjacent to DBA-associated gene *RPL35A* in the human genome. In DBA-RPL35A patients with a large 3q29 deletion containing *IQCG*, there was a 3.5-fold increase in microcephaly cases compared to other RPL35A variants (Gianferante et al. 2021). Given the role of IQCG in centrosome function, MT organization and cell division, this observation holds significant importance. Furthermore, Ninein helps maintain neural progenitor cell fate by retaining older centrosomes within human cortical organoids (Royall et al. 2023). Considering potential cooperation among IQCG-GSK3β-Ninein axis, further exploration into IQCG’s involvement in neural cells is warranted. These studies will provide novel insights and valuable cues regarding the underlying biological basis that drives human evolution.

## Materials and Methods

### Cell culture and transfection

All cell lines were sourced from the Cell Bank of Kunming Institute of Zoology, Kunming, China. HeLa, U2OS and HEK293T cells were cultured at 37°C in Dulbecco’s Modified Eagle Medium (DMEM, Gibco) supplemented with 10% fetal bovine serum. Cells were grown to 50-70% confluence for transient transfections, which were performed using Lipofectamine™ 3000 transfection reagent (Thermo Fisher Scientific) according to the manufacturer’s protocol. Analyses were conducted 48 hours post-transfection.

### Plasmid construction

Fourteen expression plasmids were created using either in-fusion cloning method or ligation with T4 DNA ligase. Primers for in-fusion cloning were designed using an online tool provided by Takara Bio (https://www.takarabio.com/services-and-support/online-tools). Detailed construction methods are described in the Supplementary Information, with plasmid names and primer sequences listed in Table S1.

### SiRNAs and shRNA constructs

SiRNAs and plasmid-expressed shRNAs in pSGU6/GFP/Neo vector were obtained from Sangon Biotech (Shanghai, China). Details of the siRNAs and shRNAs including names and target regions are provided in Table S2.

### Antibodies

Details of antibodies used in this study are listed in Table S3.

### Immunofluorescence microscopy and Western blot

For immunofluorescence microscopy, cells were fixed, blocked, and stained with specific antibodies; with procedural details are provided in the Supplementary Information. Microscopy was conducted using a Leica DMi8 or Zeiss LSM 800, with image processing and analyses performed using ImageJ (Schneider et al. 2012) or QuPath (Bankhead et al. 2017). Protein samples were prepared and analyzed via Western blot as detailed in the Supplementary Information.

### Function prediction of *IQCG*

We adopted the method used in Sun et al. (Sun et al. 2022) to infer the putative function of *IQCG*. Briefly, based on more than 8,000 RNA-Seq profilings of 53 human tissues from the Genotype-Tissue Expression (GTEx) database (GTEx Consortium 2013), we first calculated the expression correlation between *IQCG* and all expressed genes to obtain the genes that were significantly correlated with *IQCG* expression after multiple testing corrections (FDR < 0.01) and then calculated the enriched biological pathways of the obtained genes using Fisher’s exact test (FDR < 0.05).

### **I**mmunoprecipitation experiment

Lysates from HEK293T cells expressing eGFP or Flag-tag proteins were incubated with anti-GFP or anti-Flag affinity beads (Smart-Lifesciences). Elutions from the beads and proteins from the original cell lysates were next analyzed by Western blot as inputs and immunoprecipitations (IPs) with corresponding antibodies. Detailed procedures are described in the Supplementary Information.

### Microtubule regrowth experiment

U2OS cells transfected with shRNA for 48h were treated with 13.2 μM Nocodazole in DMEM for 4 hours at 37℃, and then Nocodazole was removed thoroughly by washing with phosphate-buffered saline (PBS) four times. Cells were allowed to recover in DMEM at 37℃ for 0 or 5 minutes. Cells were then fixed with 4% paraformaldehyde in PBS with 0.1% Triton X-100 for 15 min, and stained with anti-β-tubulin and anti-Pericentrin (PCNT) antibodies. Images were collected as immunofluorescent samples using a microscope.

### Microtubule co-sedimentation assay

Briefly, HeLa cells expressing eGFP or eGFP-IQCG were treated by microtubule (MT) destabilizing drug Nocodazole or stabilizing drug Docetaxol. Polymerized MTs were stabilized using tubulin PEM (PIPES-EGTA-MgCl2) buffer containing glycerol. Solubles containing free tubulins were collected, as soluables (Ss). Then polymerized MTs and their interacting proteins were collected, as pellets (Ps). Ss and Ps were subject to Western blot for β-tubulin and associated protein detection. Detailed procedures are described in the Supplementary Information.

### Sources of sequences in evolutionary analyses

Protein and cDNA sequences were retrieved from Ensembl (Harrison et al. 2024) (http://www.ensembl.org/index.html), or the National Center for Biotechnology Information (NCBI, http://www.ncbi.nlm.nih.gov). We used eight animal species for aligning IQCG amino acid sequence, including *Homo sapiens* (human)*, Pan troglodytes* (chimpanzee), *Canis lupus familiaris* (dog), *Felis catus* (cat), *Equus caballus* (horse), *Gallus gallus* (chicken), *Anolis carolinensis* (lizard) and *Danio rerio* (zebrafish), using *Drosophila melanogaster* (fruitfly) as an outgroup. Accession numbers for these sequences are listed in Table S4.

For reconstructing the ancestral coding sequence (CDS) of IQCG, eleven primate species were analyzed, including *Homo sapiens* (human), *Pan troglodytes* (chimpanzee), *Gorilla gorilla* (gorilla), *Pongo abelii* (orangutan), *Nomascus leucogenys* (gibbon), *Rhinopithecus roxellana* (golden snub-nosed monkey), *Chlorocebus sabaeus* (Vervet-AGM), *Macaca fascicularis* (macaca), *Papio anubis* (olive baboon), *Aotus nancymaae* (night monkey), *Callithrix jacchus* (marmoset). *Mus musculus* (mouse) served as the outgroup. Sequence integrity was checked manually, and CDS sequences that were error-free and encoded 443aa or larger were selected for further analyses. NCBI accession number or Ensembl ID for these sequences are listed in Table S5.

### Sequence alignment and phylogenetic analysis

Amino acid and nucleotide sequences were aligned using the MUSCLE algorithm (Edgar 2004) implemented in MEGA11 software (Tamura et al. 2021), and visualized with GeneDoc version 2.7.0. The phylogenetic trees were constructed using the maximum likelihood method in MEGA11, based on the Tamura-Nei model for CDS and the Poisson correction model for amino acid sequences. Positions containing gaps or missing data were completely excluded. Tree reliability was assessed by bootstrap analysis with 1000 pseudoreplicates.

### Ancestral sequence derivation

Ancestral sequences were deduced using the maximum likelihood method in MEGA11. The tree file used to deduce the ancestral sequence was used uniformly ((((((Human, Chimpanzee), Gorilla), Orangutan), Gibbon), (Golden snub-nosed monkey, (Vervet-AGM, (Macaca, Baboon)))), (Marmoset, Night monkey), Mouse).

### Tests of positive selection

Rates of non-synonymous (Ka) and synonymous (Ks) substitutions were calculated using KaKs_Calculator 2.0 (Wang et al. 2010) based on the Modified Yang-Nielsen method (Zhang et al. 2006). Files were prepared for analysis using ClustalX 2.0 and converted to the necessary formats with AXTConvertor.

### Sliding window analysis

Ka/Ks was calculated at 10 codons increment and the results were implemented in SWAKK (Lee et al. 2008) (https://ibl.mdanderson.org/swakk/). To avoid stochastic variation of Ks being zero, Ks of the entire gene was used as denominator.

### Database analyses of common interaction protein of IQCG

We searched for common interaction protein of IQCG in Harmonizome 3.0 database (Rouillard et al. 2016) that intersected the protein-protein interactions from low-throughput or high-throughput studies from the following database sources: Reactome, NCI Pathways, PhosphoSite, HumanCyc, HPRD, PANTHER, DIP, BioGRID, IntAct, BIND, Transfac, MiRTarBase, Drugbank, Recon X, Comparative Toxicogenomics Database, and KEGG.

### Statistical analysis

Statistical analyses were performed using GraphPad Prism 9.0. Differences were evaluated using unpaired Student’s t-test, with significance set at *p* < 0.05. Data were presented as means ± SD.

### Data Availability

The data underlying this article are available in the article and in its online Supplementary Information.

## Supplementary Information

Supplementary Information is available at https://doi.org/(to be determined).

## Acknowledgments

The authors thank Dr. Jian Zhang for sharing cell culture facilities and Dr. Peng Shi for suggestion on evolutionary analysis This work was supported by National Natural Science Foundation of China (31871257) and Science and Technology Planning Project in Key Areas of Yunnan Province (202303AP140004) to HY, National Natural Science Foundation of China (32070695) and Yunnan Province Basic Research Program (202101AW070016, 202201AT070071 and 202301BF070001-012) to WF, National Key R&D Program of China (2022YFC2303200) and National Natural Science Foundation of China (82471347) to HH.

## Author Contributions

Conceptualization and writing original draft: XL, WF, HH and HY. Experiment and analysis: XL, YW, XH, YS. All authors read and approved the final manuscript.

---

**Figure S1 - IQCG knockdown decreases total microtubule mass and impairs recruitment of centrosome proteins. (A)** The knockdown of IQCG by siRNA in HEK293T cells is confirmed through Western blot analysis. **(B)** IQCG knockdown decreased total microtubule mass in HeLa cells with shRNA expression, revealed by immunofluorescent staining. Arrows show the loss of radical MT pattern from the centrosome. Green: GFP fluorescence marking shRNA expression; red: microtubule revealed by anti-β-tubulin; blue: nucleus stained by DAPI. n ≥ 200. Scale bar: 10 µm. **(C)** IQCG knockdown impairs the recruitment of centrosome proteins Ninein, PCM1, and Rootletin, as well as the distribution of centrosomal peripheral MT minus-end protein CAMSAP2. Boxed areas are enlarged for detail. Arrows highlight reduced or vague location of the proteins at the centrosome. Gray: GFP fluorescence marking shRNA expression; green: centrosome stained by anti-PCNT; red: CAMSAP2, Ninein, PCM1 and Rootletin stained by specific antibodies; blue: nucleus stained by DAPI. Immunofluorescence staining area of these proteins was quantitatively analyzed in control or shRNA-expressing HeLa cells. n = 30. Scale bars: 10 µm; enlarged boxes 2 µm. Three independent experiment replications were performed.

**Figure S2 - IQCG interacts with tubulin regardless of the polymerization status of MTs. (A)** Treatment of HeLa cells with Nocodazole or Docetaxel depolymerizes or polymerizes MTs, respectively. The soluble (S) contains free tubulin and polymerized MT-unbound fraction, and pellet (P) contains polymerized MTs and their bound fraction. MT polymerization status is blotted by anti-β-tubulin antibody. **(B)** Co-sedimentation assays show that eGFP-IQCG’s allocation in the solubles and the pellets remains consistent (indicated by the red boxes), irrespective of the polymerization status of MTs as revealed by anti-eGFP antibody. Three independent experimental replications were performed.

**Figure S3 - Functional domains and acid-base characteristics of the human IQCG protein, and the correct expression of eGFP-IQCG and its fragments. (A)** Schematic representation of the four functional regions divided in the human IQCG protein. An IQ motif and several disordered and coiled regions are indicated. **(B)** Pie charts depict the amino acid composition of these regions, with green segments representing basic amino acids and red segments representing acidic amino acids. The proportion of acidic and basic amino acids, along with the isoelectric point (pI), are provided for each region. **(C)** Correct expression of eGFP-IQCG and its fragment constructions in HEK293T cells is confirmed by Western blot analysis using anti-eGFP antibody. All constructions are correctly expressed, except that eGFP-IQCG-M exhibited low levels due to a potentially detrimental effect on cell viability. The eGFP-IQCG-M itself was therefore not used in further co-IP experiments. Note: The N1 region displays a larger band than predicted, likely as a result of its unusually low isoelectric point.

**Figure S4 - Endogenous GSK3β localizes at the centrosome in both interphase and mitosis. (A)** Correct expression of eGFPGSK3β in HEK293T cells is confirmed by Western blot analysis using anti-eGFP antibody. **(B)** Centrosome localization of endogenous GSK3β is detected by anti-GSK3β (left), which is confirmed by transiently expressed eGFP-GSK3β fusion protein (right) in interphase HeLa cells. EGFP alone serves as a control (middle). Boxed areas are enlarged for detail. Green: endogenous GSK3β detected by anti-GSK3β, eGFP or eGFP- GSK3β fluorescence; red: centrosome stained by anti-PCNT. Scale bars: 10 µm; enlarged images 1 µm. **(C)** The knockdown of GSK3β by siRNA in HEK293T cells is confirmed by Western blot analysis. ***p < 0.001. **(D)** GSK3β remains significant at the centrosome throughout mitosis in HeLa cells. Green: endogenous GSK3β stained by anti- GSK3β; red: centrosome stained by anti-PCNT; blue: chromosome stained by DAPI. Scale bar: 10 µm. Three independent experimental replications were performed.

**Figure S5 - Protein sequence alignment of IQCG across eight vertebrate species reveals significant variability in the N2 region.** The N1, N2, M, and C regions of the IQCG protein are denoted by color-coded bars above in blue, yellow, green and red, respectively.

