## Supplementary Information for "Dual roles of IQCG, a novel microtubule nucleation factor rapidly evolving in humans"

containing:

Supplementary Figures 1-5 (Figure legends in the main text)

Supplementary Tables 1-5

Supplementary Materials and Methods

#### Supplementary Figures 1-5

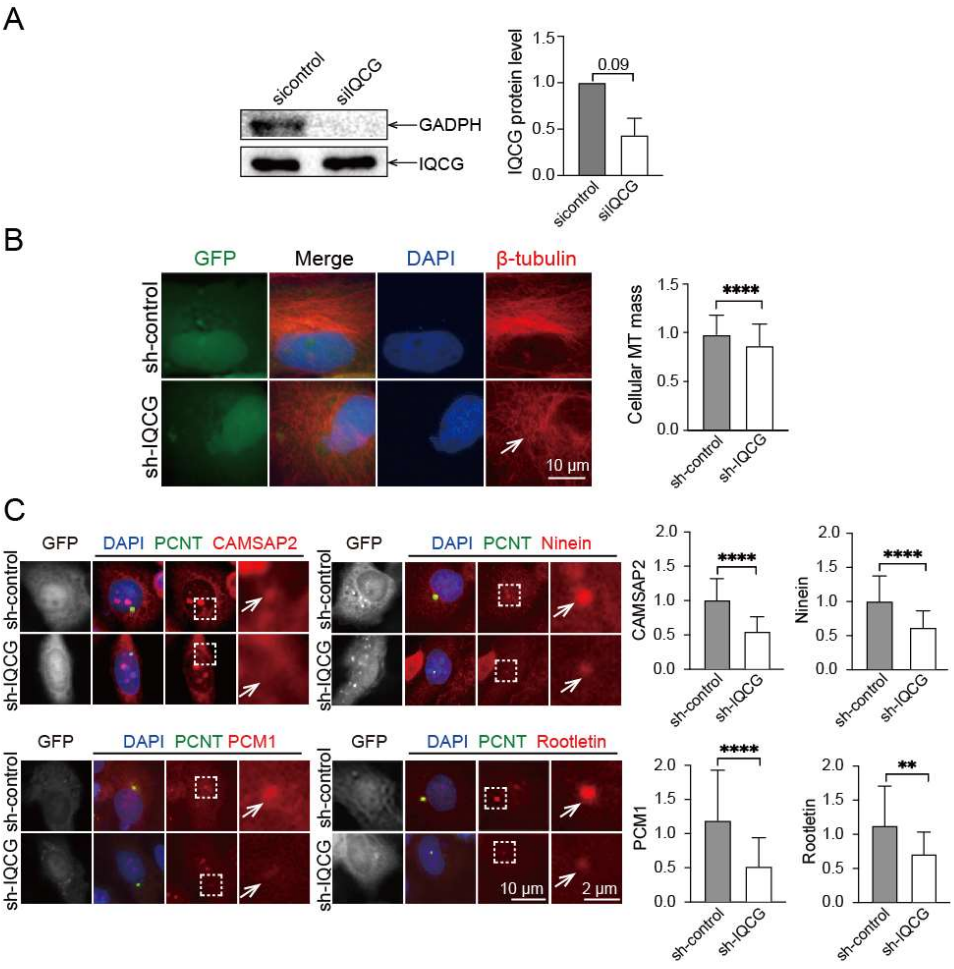


Figure S1

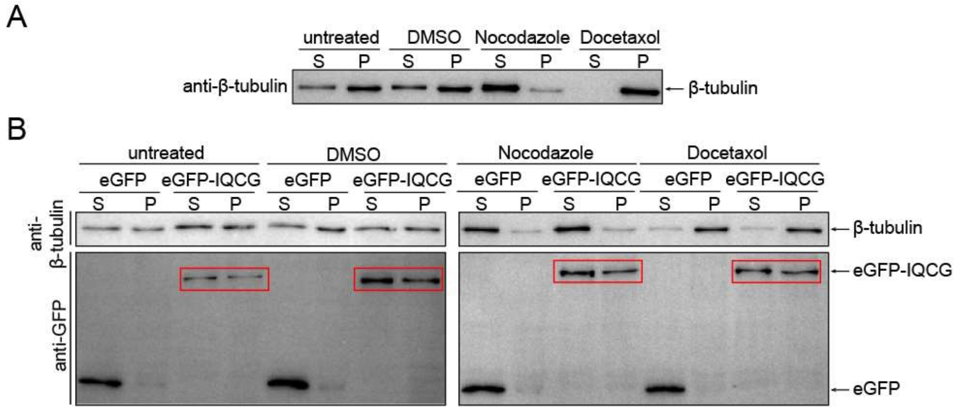
A
untreated
DMSO
Nocodazole
Docetaxol
S P S P S P S P
anti-β-tubulin
β-tubulin
B
untreated
DMSO
Nocodazole
Docetaxol
eGFP
eGFP-IQCG
S P S P
anti-β-tubulin
anti-GFP
β-tubulin
eGFP-IQCG
eGFP

1

2 Figure S2

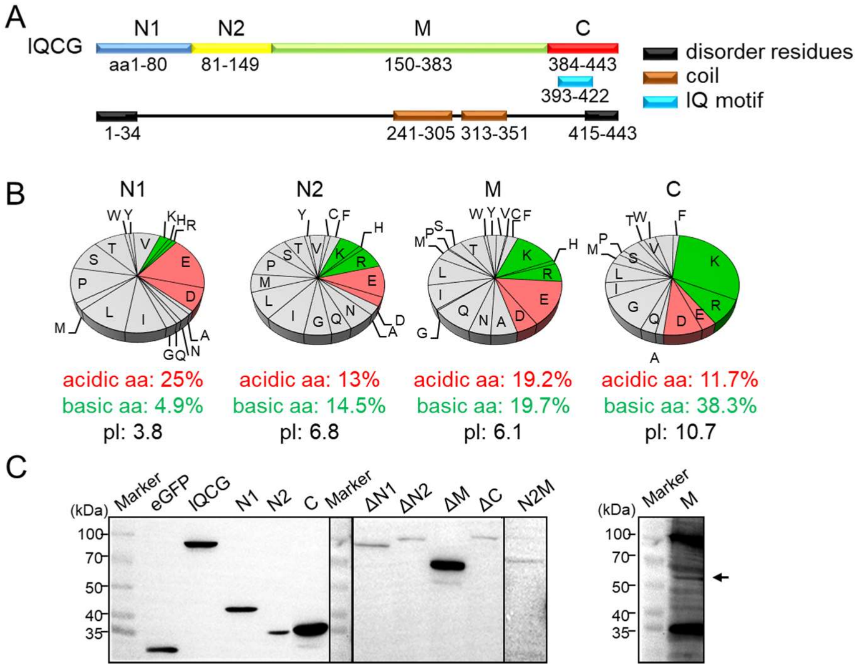
A
IQCG
N1
N2
M
C
aa1-80
81-149
150-383
384-443
393-422
1-34
241-305
313-351
415-443
disorder residues
coil
IQ motif
B
N1
N2
M
C
acidic aa: 25%
basic aa: 4.9%
pI: 3.8
acidic aa: 13%
basic aa: 14.5%
pI: 6.8
acidic aa: 19.2%
basic aa: 19.7%
pI: 6.1
acidic aa: 11.7%
basic aa: 38.3%
pI: 10.7
C
(kDa)
Marker
eGFP
IQCG
N1
N2
C
Marker
ΔN1
ΔN2
ΔM
ΔC
N2M
100
70
50
40
35
(kDa)
Marker
M
100
70
50
40
35

Figure S3

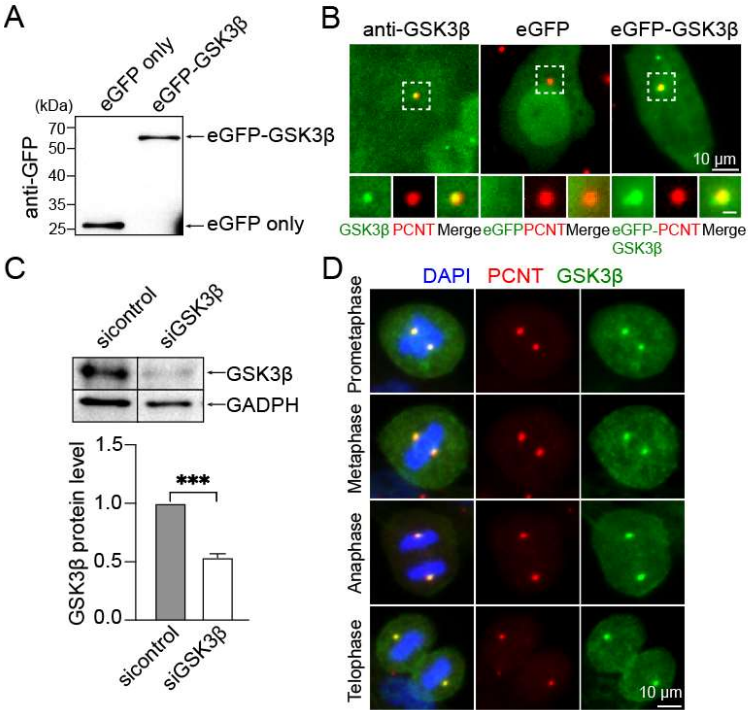
A
eGFP only
eGFP-GSK3β
(kDa)
70
50
40
35
25
anti-GFP
eGFP-GSK3β
eGFP only
B
anti-GSK3β
eGFP
eGFP-GSK3β
10 μm
GSK3β
PCNT
Merge
eGFP
PCNT
Merge
eGFP-GSK3β
PCNT
Merge
C
sicontrol
siGSK3β
GSK3β
GADPH
GSK3β protein level
1.5
1.0
0.5
0.0
***
sicontrol
siGSK3β
D
DAPI
PCNT
GSK3β
Prometaphase
Metaphase
Anaphase
Telophase
10 μm

Figure S4

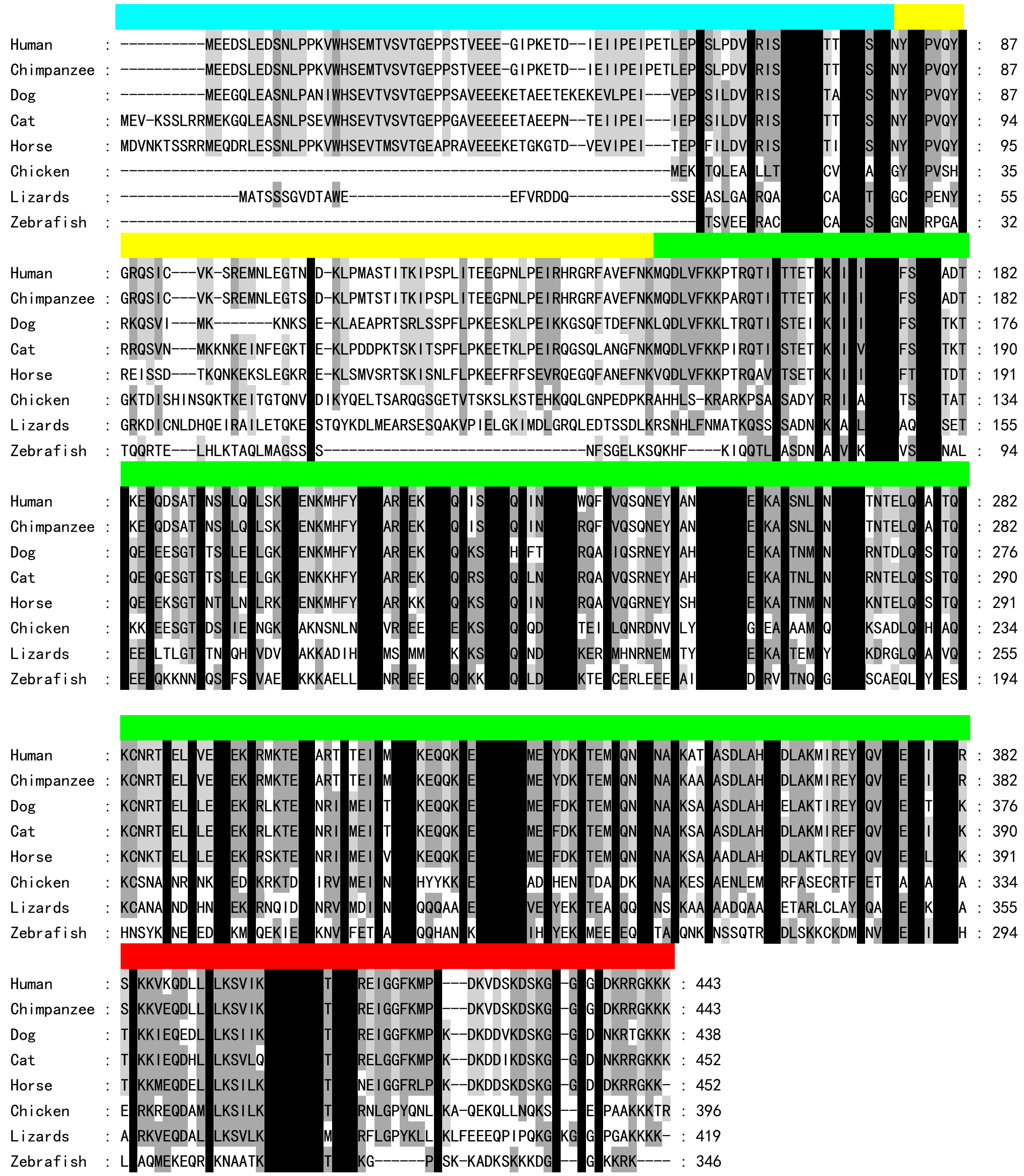

Figure S5

### Supplementary Tables 1-5

**Table S1. List of plasmid names and primers used to construct plasmids.**

| Plasmid | PCR primer | Primer sequence |
|---|---|---|
| eGFP-GSK3β | eGFP-GSK3β-U | GGACTCAGATCTCGAATGTCAGGGCGGCCCAGAACCA |
| | eGFP-GSK3β-L | TAGATCCGGTGGATCCTCAGGTGGAGTTGGAAGCTGAT |
| eGFP-IQCG | eGFP-IQCG-U | GGACTCAGATCTCGAATGGAAGAAGACAGCCTG |
| | eGFP-IQCG-L | TAGATCCGGTGGATCCTCACTTCTTCTTGCCTCTC |
| eGFP-IQCG - N1 | eGFP-N1-U | GGACTCAGATCTCGAATGGAAGAAGACAGCCTG |
| | eGFP-N1-L | TAGATCCGGTGGATCCGTTAGATCCGGTGGATCCGT |
| eGFP-IQCG - N2 | eGFP-N2-U | GGACTCAGATCTCGAATCATGCCCGTTCAGT |
| | eGFP-N2-L | TAGATCCGGTGGATCCAAGATCCTGCATTTTGTTA |
| eGFP-IQCG - M | eGFP-M-U | GGACTCAGATCTCGAGTCTTCAAAAAACCTACA |
| | eGFP-M-L | TAGATCCGGTGGATCCGCTCCTCTCCTTTTCTATA |
| eGFP-IQCG - C | eGFP-C-U | GGACTCAGATCTCGAAAGAAGAAGGTAAAACAGG |
| | eGFP-IQCG-L | TAGATCCGGTGGATCCTCACTTCTTCTTGCCTCTC |
| eGFP-IQCG - N1N2 | eGFP-N1-U | GGACTCAGATCTCGAATGGAAGAAGACAGCCTG |
| | eGFP-N2-L | TAGATCCGGTGGATCCAAGATCCTGCATTTTGTTA |
| eGFP-IQCG - N2M | eGFP-N2-U | GGACTCAGATCTCGAATCATGCCCGTTCAGT |
| | eGFP-M-L | TAGATCCGGTGGATCCGCTCCTCTCCTTTTCTATA |
| eGFP-IQCG - MC | eGFP-M-U | GGACTCAGATCTCGAGTCTTCAAAAAACCTACA |
| | eGFP-IQCG-L | TAGATCCGGTGGATCCTCACTTCTTCTTGCCTCTC |
| eGFP-IQCG-ΔN1 | eGFP-IQCG-ΔN1-U | GGACTCAGATCTCGAATCATGCCCGTTCAGT |
| | eGFP-IQCG-ΔN1-L | TAGATCCGGTGGATCCTCATCACTTCTTCTTGCCTCTC |
| *eGFP-IQCG-ΔN2* | *eGFP-IQCG-M-U* | *CGCGGATCCGTCTTCAAAAAACCTACA* |
| | *eGFP-IQCG-C-L* | *CGCGGATCCTCACTTCTTCTTGCCTCTC* |
| *eGFP-IQCG-ΔM* | *eGFP-IQCG-C-U* | *CGCGGATCCAAGAAGAAGGTAAAACAGG* |
| | *eGFP-IQCG-C-L* | *CGCGGATCCTCACTTCTTCTTGCCTCTC* |
| eGFP-IQCG-ΔC | eGFP-N1-U | GGACTCAGATCTCGAATGGAAGAAGACAGCCTG |
| | eGFP-M-L | TAGATCCGGTGGATCCGCTCCT CTCCTTTTCTATA |
| Flag-tubulin | Flag-tubulin-U | CGTCAGATCCGCTAGCGCTAGCGCTACCGGACT |
| | Flag-tubulin-L | CACGCACTCGAGATCTACGAATTCGAAGCTTGAGC |

Note: Plasmids in italic are constructed by T4 ligation method; all others are constructed by in-fusion cloning method.

**Table S2. List of names and target sites of siRNA and shRNA.**

| Knockdown RNA | Target position in CDS (bp) | Target sequence（5'-3'） |
|---|---|---|
| siIQCG#1 and sh-IQCG#1 | IQCG 101-123 | AAGAAGGAATACCTAAAGAAACA |
| siIQCG#3 and sh-IQCG#3 | IQCG 1242-1264 | TGGTTTCAAGATGCCTAAAGACA |
| siGSK3β | GSK3β 855-878 | CCCAAACTACACAGAATTTAATT |

Note: SiIQCG#1 and #3, or sh-IQCG#1 and #3, are used in combination in knockdown experiments. SiIQCG#1 and sh-IQCG#1, or siIQCG #3 and sh-IQCG #3, target the same region, respectively.

**Table S3. List of primary and secondary antibodies used in immunofluorescence microscopy and Western blot experiments.**

| Application | Antibody | Producer | Catalog number | Source | Dilution |
|---|---|---|---|---|---|
| Immunofluorescent primary antibody | anti-IQCG | CUSABIO | CSB-PA867116LA01HU | rabbit | 1:200 |
| | anti-β-tubulin | Proteintech | 66240-I-Ig | mouse | 1:200 |
| | anti-CP110 | Proteintech | 12780-I-AP | rabbit | 1:250 |
| | anti-CAMSAP2 | Proteintech | 178880-1-AP | rabbit | 1:200 |
| | anti-PCNT | Abcam | ab4448 | rabbit | 1:200 |
| | anti-PCNT | Abcam | ab28144 | mouse | 1:300 |
| | anti-β-tubulin | Abcam | ab21057 | goat | 1:500 |
| | anti-Ninein | Santa Cruz | sc-376420 | mouse | 1:200 |
| | anti-Rootletin | Santa Cruz | sc-374056 | mouse | 1:200 |
| | anti-PCM1 | Santa Cruz | sc398366 | mouse | 1:200 |
| | anti-GSK3β | Huaan | ET1607-71 | rabbit | 1:400 |
| | anti-Centrin | Merck Millipore | 04-1624 | mouse | 1:300 |
| Immunofluorescent secondary antibody | Alexa Fluor 488-conjugated donkey anti-mouse | YEASEN | 34106ES60 | donkey | 1:200 |
| | Alexa Fluor 488-conjugated goat anti-mouse | Beyotime | A0428 | goat | 1:200 |
| | Alexa Fluor 488-conjugated goat anti-rabbit | Beyotime | A0423 | goat | 1:200 |
| | Alexa Fluor 647-conjugated donkey anti-rabbit | YEASEN | 34213ES60 | donkey | 1:200 |
| | Alexa Fluor 647-conjugated donkey anti-mouse | YEASEN | 34106ES60 | donkey | 1:200 |
| | Cy3-conjugated donkey anti-goat | Beyotime | A0502 | donkey | 1:200 |
| Western blot primary antibody | anti-GFP | Sangon | NB100-1614 | mouse | 1:500 |
| | anti-Flag | Novusbio | NB600-344 | goat | 1:500 |
| | anti-GSK3β | Cell Signaling Technology | 12456 | rabbit | 1:2000 |
| | anti-IQCG | CUSABIO | CSB-PA867116LA01HU | rabbit | 1:8000 |
| | anti-β-tubulin | Proteintech | 66240-I-Ig | mouse | 1:8000 |
| Western blot secondary antibody | goat anti-rabbit HRP | Beyotime | A0208 | goat | 1:5000 |
| | donkey anti-goat HRP | Beyotime | A0181 | donkey | 1:5000 |
| | goat anti-mouse HRP | Beyotime | A0216 | goat | 1:5000 |

**Table S4. List of NCBI accession numbers of IQCG proteins from nine animal species.**

| Species | NCBI accession number | Length (aa) |
|---|---|---|
| Human | NP_001127907.1 | 443 |
| Chimpanzee | XP_054538516.1 | 443 |
| Dog | XP_005639646.1 | 438 |
| Cat | XP_006936192.2 | 452 |
| Horse | XP_014588325.1 | 452 |
| Chicken | XP_040534634.1 | 409 |
| Lizard | XP_003227015.1 | 419 |
| Zebrafish | NP_001333169.1 | 346 |
| Fruit fly | NP_651566.1 | 448 |

**Table S5. List of NCBI accession numbers or Ensembl IDs of IQCG transcripts from 11 primates and one outgroup species.**

| Species | Transcript ID |
|---|---|
| Human | NM_032263.5 |
| Chimpanzee | XM_054682541 |
| Gorilla | ENSGGOT00000048708.1 |
| Orangutan | XM_024241189.3 |
| Gibbon | XM_030823061.1 |
| Golden snub-nosed monkey | XM_017865486.1 |
| Vervet-AGM | ENSCSAT00000006500.1 |
| Macaca | XM_015132645 |
| Olive baboon | XM_009198831.4 |
| Marmoset | XM_017966000.3 |
| Night monkey | XM_012476623.2 |
| Mouse | XM_011246004.4 |

### Supplementary Materials and Methods

**Expression plasmid construction**

A total of 14 expression plasmids were constructed in this study as follows. For the eGFP-series expression plasmids, pEGFP-C1-Tubulin (Miaoling Biology) was used as a vector. Full-length coding sequences of IQCG, GSK3β, or IQCG fragments including N1, N2, M, C, N1N2, N2M, MC, ΔN1 and ΔC inserts were PCR-amplified from human cDNA and in-fusion cloned (ClonExpress II One Step Cloning Kit, Vazyme) into the large fragment resulted from XhoI-BamHI (New England Biolabs) cut of pEGFP-C1-Tubulin, replacing the tubulin fragment and omitting the XhoI site. To generate the eGFP-ΔM plasmid, the IQCG-C fragment was PCR-amplified, digested with BamHI, and ligated into the BamHI site of previously constructed pEGFP-C1-N1N2 using T4 ligase (New England Biolabs). Similarly, the eGFP-ΔN2 plasmid was constructed by inserting the PCR-amplified and digested IQCG-MC fragment into the BamHI site of pEGFP-C1-N1 using T4 ligase.

In constructing pFlag-C1-Tubulin, the flag fragment was PCR-amplified and then in-fusion cloned with the large fragment of pEGFP-C1-Tubulin linearized by NheI and BglII restriction enzymes (New England Biolabs).

**Immunofluorescence microscopy**

HeLa cells were cultured in 24-well plates for wide-field microscopy or 4-chamber glass bottom dishes for confocal microscopy. Cells were fixed with pre-cooled (-20°C) methanol for 5 minutes or 4% formaldehyde (room temperature) for 10 minutes, followed by blocking in buffer containing 0.5% Tween-20 and 10% blockaid solution (Thermo Fisher Scientific) in PBS for 1 hour. Cells were then incubated sequentially with primary and secondary antibodies for 1 hour each at room temperature and visualized by microscopes.

**Western blot**

Cells were lysed in RIPA buffer (Beyotime) supplemented with 5 mM PMSF (Beyotime), then sonicated and centrifuged at 4°C for 10 minutes at 12,000 rpm. Protein concentrations were determined using the BCA method (Biosharp). Equal protein amounts were separated by 8-10% SDS-PAGE under reducing conditions and transferred to PVDF membranes. The membranes were blocked with skim milk for 1 hour and incubated overnight at 4°C with primary antibodies in the dilution solution (Solarbio). After washing in PBST, the membranes were incubated with HRP-conjugated secondary antibodies for 90 minutes at room temperature. Immunoreactive bands were detected using ECL reagent (Biosharp). Images were analyzed using ImageJ software.

**Co-immunoprecipitation experiment**

HEK293T cells were collected 48 hours after transfection, washed in PBS and lysed on 4 ℃ for one hour in lysis buffer A [50 mM HEPES, 0.3 mM CHAPS, 1% Protease Inhibitor Cocktail (Bimake)]. After centrifugation at 13,000 rpm for 10 min at 4 ℃, 1/10 of the lysate was retained for input or total cell lysate analysis by Western blot. Immunoprecipitations were performed by incubating the lysate with 20 μl of anti-GFP or anti-Flag affinity Beads (Smart-Lifesciences) for 3 hours at 4°C with slow rotation. Beads were washed three times in buffer B (50 mM HEPES, 120 mM NaCl, 0.3 mM CHAPS) at 4°C and eluted in 1× SDS loading buffer at 95°C for 5 minutes for SDS-PAGE analysis.

**Microtubule co-sedimentation assay**

HeLa cells expressing eGFP or eGFP-IQCG were cultured in 6-well plates and treated with microtubule-destabilizing drug Nocodazole (Beyotime, 0.8 μg/ml) or stabilizing drug Docetaxol (Beyotime, 5 μg/ml) for 4 hours. Cells were then incubated with 320 μl of PEM solution (1 M $MgCl_2$, 10 mM EGTA, 1M pH=7 PIPES, 0.5% Triton X-100, and 2.5% glycerol) for 2 minutes at 37°C. The solubles containing free tubulin were collected while the cell remanants were lysed in RIPA buffer, centifuged and collected as pellets, which contained polymerized tubulin and associated proteins. Solubles and pellets were analyzed by Western blot to detect microtubule-associated proteins.
